# Improved deterrence of birds using an artificial predator, the RobotFalcon

**DOI:** 10.1101/2022.05.18.492297

**Authors:** Rolf F. Storms, Claudio Carere, Robert Musters, Hans van Gasteren, Simon Verhulst, Charlotte K. Hemelrijk

**Author notes:** **Corresponding author** Rolf F. Storms, MSc, Groningen Institute for Evolutionary Life Sciences, University of Groningen, Nijenborg 7, 9747 AG Groningen, The Netherlands.

## Abstract

Collisions between birds and airplanes, bird strikes, can damage aircrafts, resulting in delays and cancellation of flights, costing the international civil aviation industry more than 1.4 billion U.S. dollars annually. Bird deterrence is therefore crucial, but the effectiveness of all available deterrence methods is limited. For example, live avian predators can be a highly effective deterrent, because potential prey will not habituate to them, but live predators cannot be controlled with sufficient precision. Thus, there is an urgent need for new deterrence methods. To this end we developed the RobotFalcon, a device that we modelled after the peregrine falcon, a cosmopolitan predator that preys on a large range of bird species. Mimicking natural hunting behaviour, we tested the effectiveness of the RobotFalcon to deter flocks of corvids, gulls, starlings and lapwings. We compared its effectiveness with that of a drone, and of conventional methods routinely applied at a military airbase. We show that the RobotFalcon scared away bird flocks from fields immediately, and these fields subsequently remained free of bird flocks for hours. The RobotFalcon outperformed the drone and the best conventional method at the airbase (distress calls). Importantly, there was no evidence that bird flocks habituated to the RobotFalcon. We propose the RobotFalcon to be a practical and ethical solution to drive away bird flocks with all advantages of live predators but without their limitations.

**Highlights:**

- We present and test a new method of deterring of deterring birds, the RobotFalcon.
- The RobotFalcon chased away flocks fast and prevented early returns.
- The RobotFalcon outperformed both a drone and convential methods.
- No evidence of habituation to the RobotFalcon was found during the study period.

## 1. Introduction

Bird strikes, birds colliding with aircrafts, cost the civil aviation industry more than 1.4 billion U.S. dollars annually (DeVeault et al., 2017; FAA, 2016b; Dolbeer et al., 2014), and in the last century bird strikes have lead to over 450 deaths in military aviation alone (Thorpe, 2016; Richardson & West, 2000; Pfeiffer et al., 2018). In agriculture, gregarious birds eating crops cause economic damage, and bird flocks can cause discomfort in urban environments (Johnson and Glahn, 1997; Thearle, 2013). Thus, besides the joy birds give to many, flocks of birds can cause non-negligble economic loss and safety hazards (Conover, 2002).

To mitigate these societal costs, there is a need to deter birds from specific locations. Many ways to deal with such human-bird conflicts have been explored. Habitats have been made unattractive to birds, but this by itself seldom solves the problem (DeVault et al., 2013; ACI, 2005), creating the need for deterrence methods. Some methods actively harm animals, e.g. taking away their eggs (Harris & Davis, 1998), blinding them with a laser (Blackwell et al., 2002), trapping and releasing them remotely, or even killing them (live shooting and falconry, Harris & Davis, 1998). Other methods rely on acoustic (pyrotechnics and distress calls) or visual deterrents (dogs, falcon silhouettes, and scarecrows) (Bishop et al., 2003; Harris & Davis, 1998). No method can clear areas from birds indefinitely and the time until birds return varies per method. Many methods suffer from varying degrees of habituation: after repeated exposure, birds respond less (Blumstein, 2016). Given the variable and temporary effectiveness of available methods there is an urgent need for new and more effective methods.

Habituation is reduced when methods resemble natural threats, such as falconry (Harris & Davis, 1998; Cook et al, 2008; Raderschall et al., 2011). However, breeding and training falcons is very costly, and its effectiveness is limited because falcons cannot be flown often and guiding their attacks is problematic (MacKinnon, 2004; Harris & Davis, 1998). Instead of live falcons, models that mimic predators may be a promising way to deter birds (e.g. Egan et al, 2020), with the advantages of a live predator, but with fewer practical limitations. We therefore developed an artificial raptor, the RobotFalcon, inspired specifically by a peregrine falcon (*Falco peregrines*), because the peregrine falcon hunts a wide spectrum of bird species over a large part of the globe (e.g. Zoratto et al., 2010; Ponitz et al., 2014). The RobotFalcon closely resembles the peregrine falcon in its shape, the coloration of its wings, beak, and head, and its relative dimensions of wings and tail (Figure 2a). It has the advantanges that it can be precisely steered to target a flock and can be flown more frequently than live falcons. Steering from the RobotFalcon is done from its perspective via a camera on its back (Figure 2b, First Person View).

**Figure 1.**
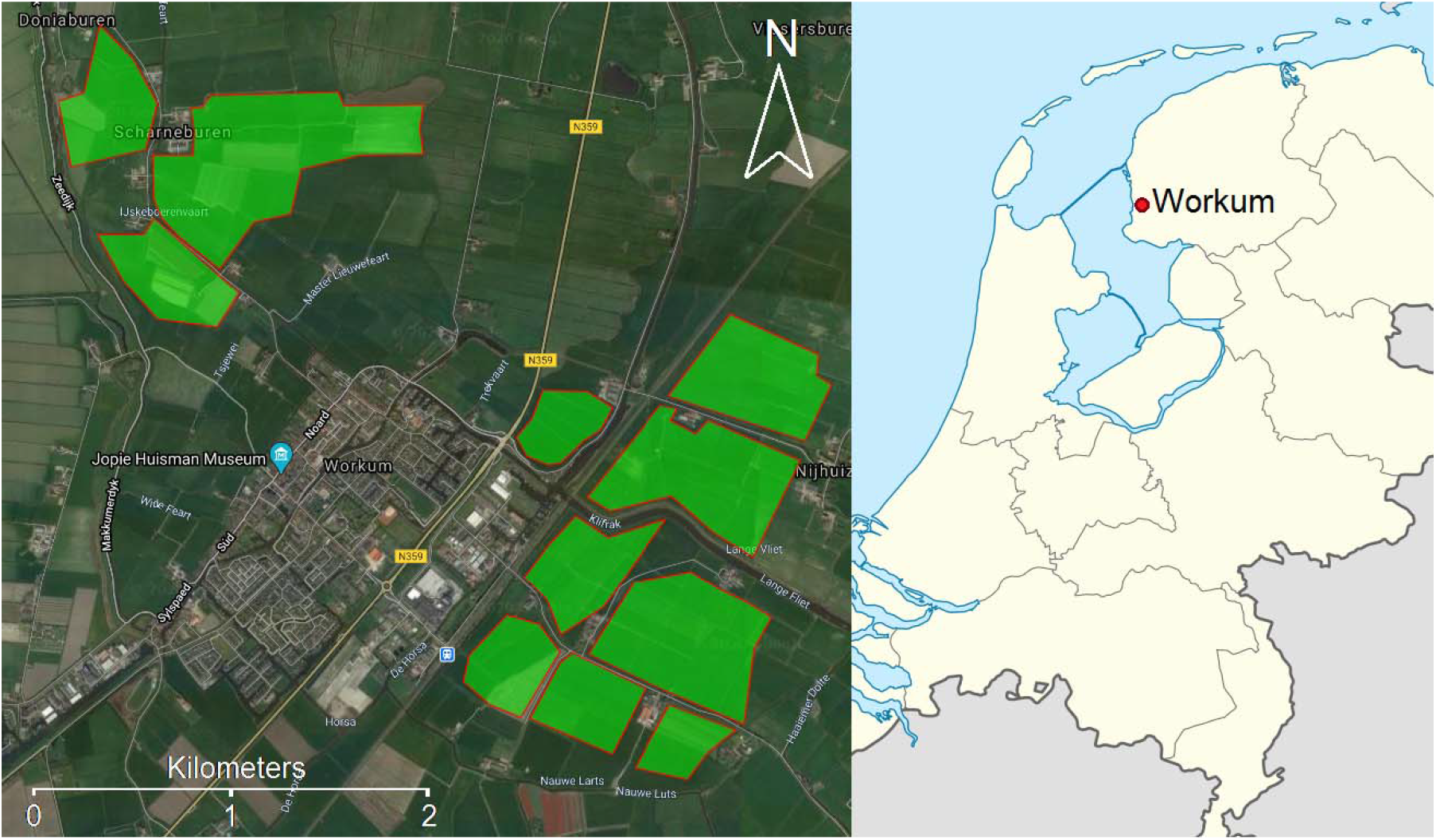
The research fields used for field experiments in Workum, highlighted in green.

**Figure 2.**
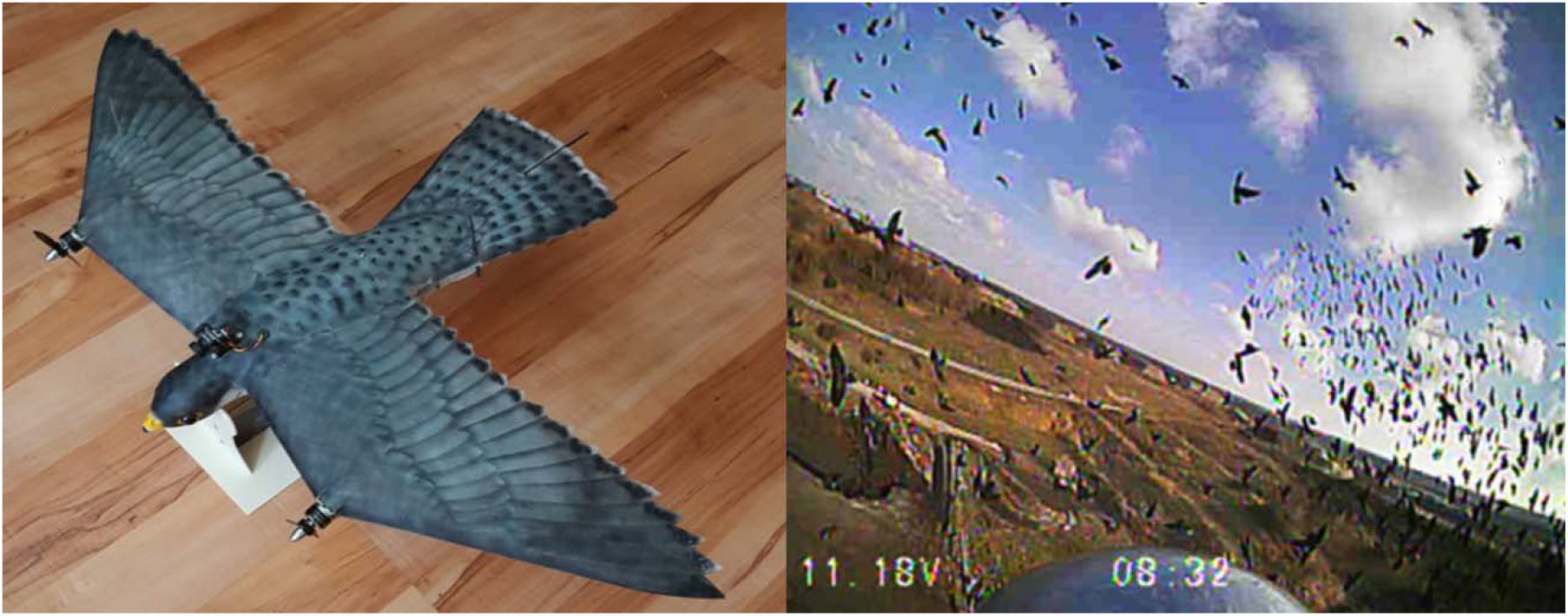
The RobotFalcon (left) and an example of its view during flight (right).

We tested the effectiveness of the RobotFalcon to drive away bird flocks by measuring the proportion of flocks it drives away, how fast fields were cleared from bird flocks, how long it took for bird flocks to return, and whether birds habituated. To this end, the RobotFalcon was flown on several species in an agrarian environment (Workum, The Netherlands). Experimental comparisons were made between the behaviour of the bird flock following attacks by the RobotFalcon and control sessions without the RobotFalcon. We further compared the effectiveness of the RobotFalcon to that of a conventional drone and methods in current use at a military airport.

## 2. Materials and Methods

### 2.1 Study area

The field work was carried out at 11 locations, in the agricultural area surrounding Workum, The Netherlands (52°59’N-5°27’E). We focused our RobotFalcon hunts on corvids (*Corvus monedula, Corvus frugilegus*, and *Corvus corone*), gulls (*Chroicocephalus ridibundus* and *Larus canus*), Northern Lapwings (*Vanellus vanellus*), and starlings (*Sturnus vulgaris*). These species are common in the study area and frequently involved in conflicts with human activities and flight safety on aerodromes (MacKinnon, 2004).

### 2.2 RobotFalcon and drone

The RobotFalcon is an ornithopter developed by RM with the features of a male peregrine falcon, *Falco peregrinus* (Figure 2). The model weighs 0.245 kg, has a cruise speed of 15 m/s and includes a camera (Runcam micro swift2, 30 fps) positioned on its head enabling first-person view. It is equipped with a GPS (Marshall RT GPS), a 2s 950mah lipo-battery and could sustain flights for up to 15 minutes. The model has a wingspan of 70 cm and has two propellers, one on each wing, used for steering. The wings don’t flap. Two certified operators (RM & RW) piloted the RobotFalcon alternatingly.

A DJI Mavic Pro drone lacking any raptor features was used for comparison. The drone was black, weighed 0.734 kg, and had a maximum speed of 18 m/s (Fig. S1).

### 2.3 Field procedure

Field work was done on 34 days between February 1^st^ and November 22^nd^ 2019 by a team of three: a pilot, an operator of a ground camera (Sony FDR-AX53 4K Camcorder, 50 fps) and a coordinator with audio recorder and GPS receiver. Wind speed and direction were measured using an anemometer (Kaindl windmaster 2) and a compass (Compass Galaxy), and the cloudiness (cloudy, partially cloudy and sunny) was scored immediately prior to the flights. We avoided rain and strong wind (> 6 on the Beaufort scale). We further recorded which birds were present (species and number), their behaviour (foraging, resting or restless) and location (using a Bushnell Tour V4 Range Finder and the ground camera).

When flocks of corvids, gulls, lapwings, pigeons or starlings were spotted on the ground within the study area, a deterrence experiment started. A subset of these experiments was randomly assigned to start with a control trial. During these trials birds were monitored for ten minutes without performing any deterrence action to see whether and when the birds fly away.

If birds remained after a control or if no control trial was assigned, a deterrence action was performed with randomly either the RobotFalcon or the drone. The pilot flew the RobotFalcon or the drone such that it approached the birds in a straight line at a constant altitude. The behaviour of the birds was recorded with the ground camera and audio recordings. The flight initiation of the flock was defined as the moment when at least one individual started taking off (i.e. from the moment it started flapping its wings) and was followed by the rest of the flock.

Once the birds were airborne, the RobotFalcon or the drone pursued the flock, simulating hunting behaviour of real peregrine falcons, based on videorecordings and analyses from previous work (Zoratto et al., 2010; Storms et al., 2019, Table 1). Such simulations included pursuing and intermittent attacks.

A hunting sequence was considered successful and ended when the birds vanished from view (using 8 × 40 binocular). The RobotFalcon or the drone was subsequently landed, and this action was filmed in order to synchronize the GPS data with the footage of the ground camera. Next we monitored the field at intervals of30 minutes for up to 120 minutes in order to record the time birds returned.

### 2.4 Data extraction and statistical analysis

Footage of the ground camera was synchronized with the GPS data of the RobotFalcon using Adobe Premiere Pro, and analyzed on a frame by frame basis, recording the escape of the flocks. Deterrence success was quantified in two ways, firstly, by the proportion of deterrence actions that cleared fields from bird flocks and, secondly, by the duration the fields stayed clear of bird flocks after deterrence. The latitude, longitude and altitude of the position of the RobotFalcon at the beginning of the flight response were used to estimate the distance between the flock of birds (using the location of the flockmember closest to the RobotFalcon) and the RobotFalcon: the Flight Initiation Distance (FID). Since the drone needed to approach flocks several times before they took flight, we did not measure the FID of a flock to the drone.

Effectiveness of the RobotFalcon was compared to that of the drone and of deterrence methods applied at military airbase Leeuwarden. Deterrence data from airbase Leeuwarden were collected from 2001 till 2016 and involved methods such as bioacoustics and pyrotechnics. The proportion of deterrence actions that resulted in clearing the field of birds was compared between the RobotFalcon and drone using a two-way ANOVA. A survival analysis was performed on the time it took birds to return, accounting for instances in which the birds had not yet returned at the last visit.

We used General Linear Models for testing whether the FID changed over time, taking into account the species, the approach altitude of the RobotFalcon and weather conditions as fixed effects, and flight identity as random effect to account for the non-independence of data on multiple species deterred during a flight.

## 3. Results

All flocks were successfully deterred by the RobotFalcon within five minutes after it started its flight, with 50% of flights resulting in fields being free of birds within 70 seconds (54 flocks, Figure 3a). This contrasts strongly with the control sessions, without deterrence, in which 15% of locations were free of birds after 5 minutes (26 flocks, Figure 3a).

**Figure 3.**
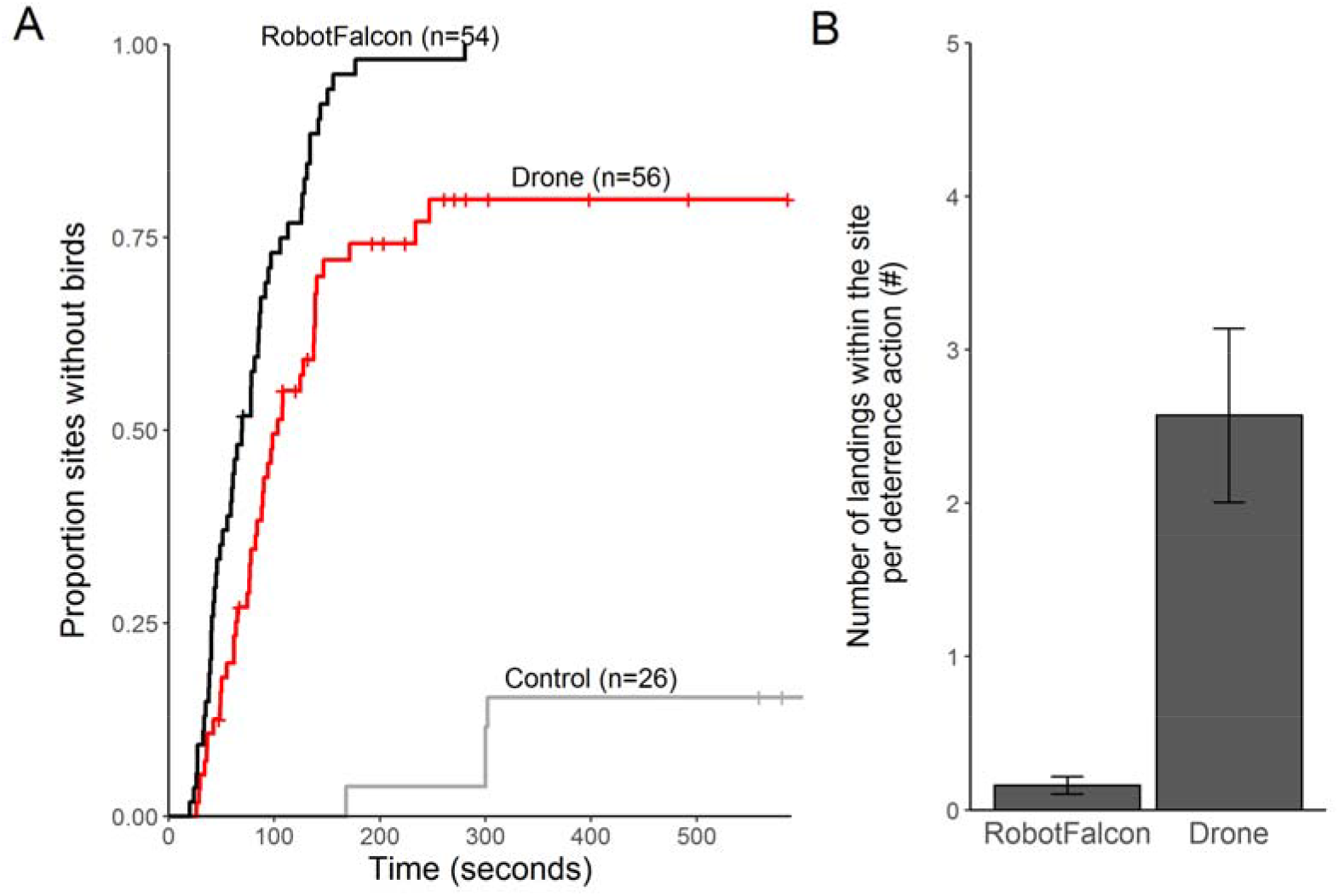
Flock responses to experimental and control flights (=no disturbance). (A). Proportion of fields cleared from flocks of birds over time after being approached by the RobotFalcon, drone or neither (control session). The three methods differed significantly (χ2(2, N =136) = 70.7, *p* < .001). (B). The average number of times flock members landed again after flying up for the RobotFalcon and drone (±SEM). Flocks landed again at the field significantly more often after flying up for the drone than the RobotFalcon (*t*(56) = 4.23, *p* < 0.01).

With the drone, it took longer to chase away flocks, and fewer fields were cleared: half of the them were cleared after 100 seconds and 80% after 5 minutes (56 flocks, Figure 3a). The RobotFalcon was also more effective in keeping flocks airborne than the drone: flocks occasionally landed briefly after taking flight, but this occurred less often when using the RobotFalcon than the drone (landing 0.2 times per hunt with the RobotFalcon versus 2.6 times with the drone; Figure 3b).

When analysed separately per species, we show that the RobotFalcon chased away flocks of corvids and gulls significantly faster than the drone, while starlings were chased away by both methods equally fast (Figure s2). Compared to the drone, flocks of all species displayed more often patterns of collective escape when hunted by the RobotFalcon (Figure 4).

**Figure 4.**
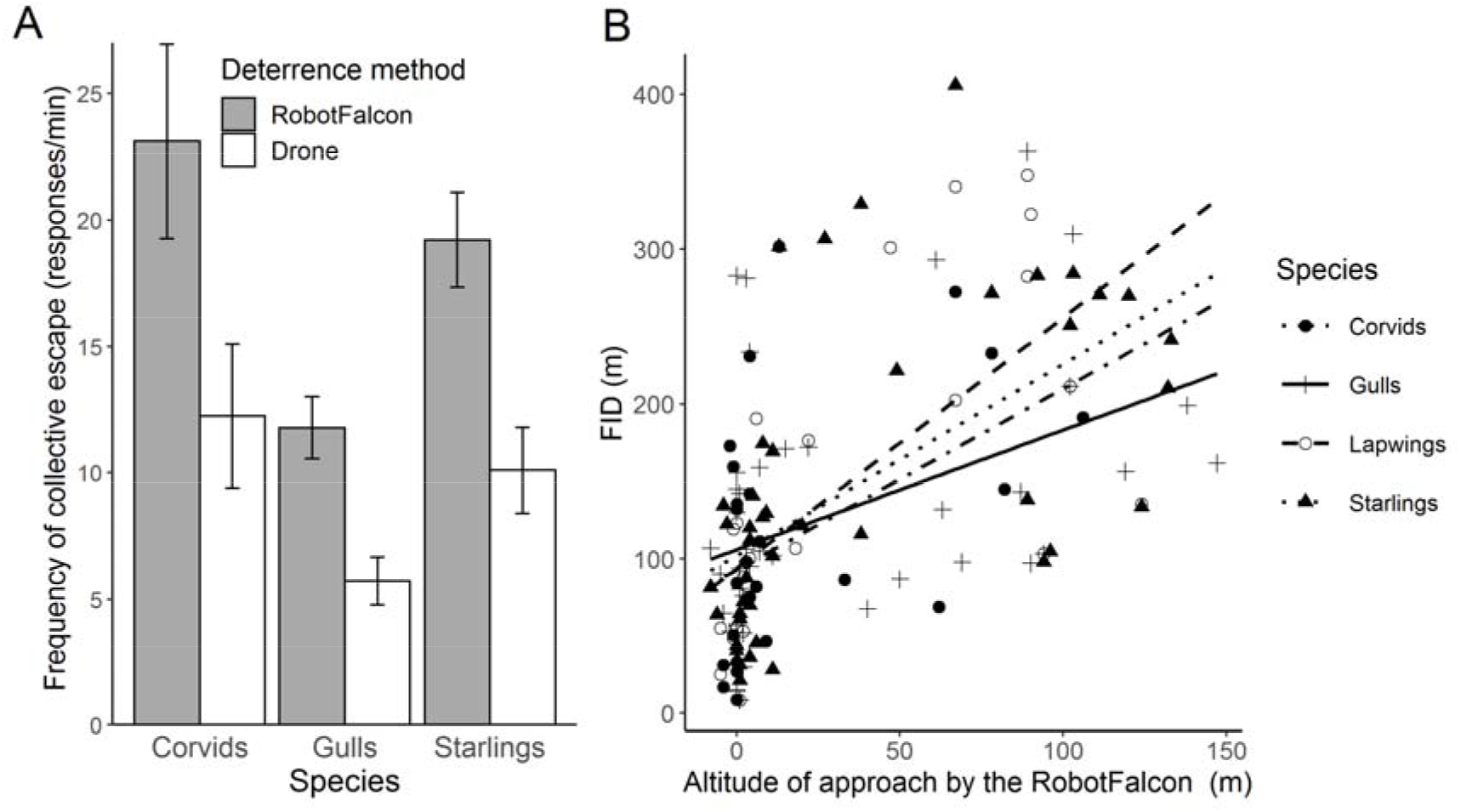
Collective escape of flocks of corvids, gulls and starlings when chased with artificial methods. A). The frequency of collective escape for the RobotFalcon and the drone (±SEM). B). The higher the approach altitude of the RobotFalcon, the further the distance at which flocks initiated flight (Flight Initiation Distance, FID).

To optimise the strategy of hunting, we studied several altitudes at which the RoborFalcon approached the flocks, and found that this affected the flight initiation distance: when the RobotFalcon approached from a higher altitude, flocks of all species fled sooner (Figure 4b).

We found no evidence for habituation to the RobotFalcon during the study period: the success at clearing fields remained high and the flight initiation distance did not change over the course of our fieldwork for any of the species (Figure 5).

**Figure 5.**
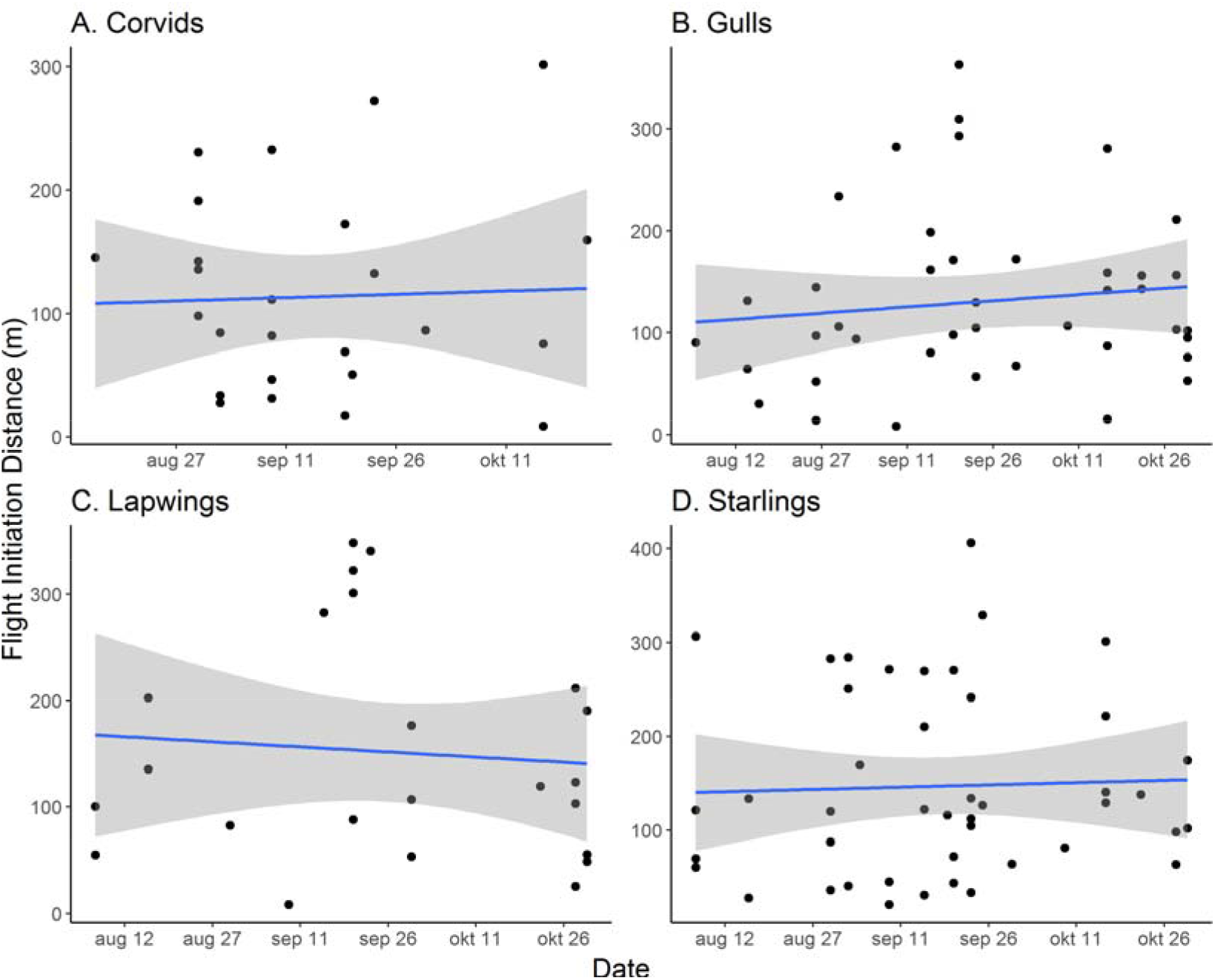
No change in the distance at which birds flocks initiated flight in response to the RobotFalcon over the period of three months of fieldwork in Workum, the Netherlands. Habituation would have resulted in a decrease of flight initiation distance over time.

Compared to the method that cleared fields of birds for the longest period at airbase Leeuwarden (the sound installation emitting calls), the RobotFalcon caused flocks of gulls, lapwings and starlings to stay away longer (Figure 6). More specifically, following flights with the RobotFalcon, flocks of starlings and lapwings stayed away for a median time of 4 hours, compared to 1.83 and 1.1 hours, respectively when deterred by distress calls. Flocks of gulls stayed away for a median time of 3 hours after flights with the RobotFalcon versus 1.5 hours when scared by distress calls. Corvids stayed away equally long when deterred by the RobotFalcon and distress calls (about an hour for both methods, Figure 6).

**Figure 6.**
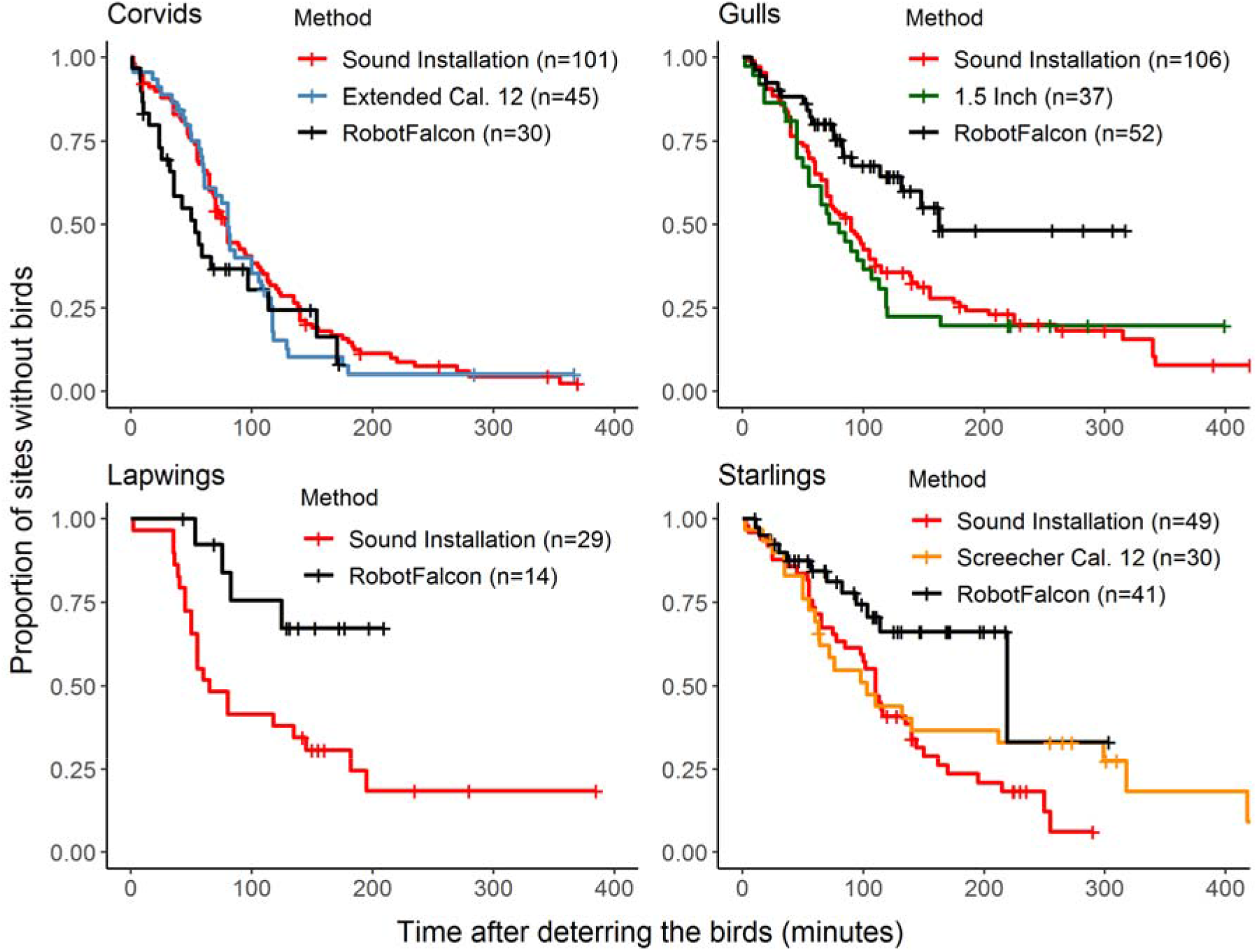
Proportion of fields without birds after deterrence with the RobotFalcon or other methods. Proportion of fields that was without birds over time after flocks of corvids, gulls, lapwings and starlings were chased away. For the airbase Leeuwarden, we show only the results for the method with the best results for each species. The sound installation method involves playing back distress calls of the species under concern. The Extended Cal. 12, 1.5 Inch and Screecher Cal. 12 are all variants of pyrotechnics. Gulls, lapwings and starlings stayed away significantly longer when chased away with the RobotFalcon than with distress calls (χ2(2, N =195) = 10.4, p = .006 ;χ2(1, N =43) = 5.9, p = .02; χ2(2, N =120) = 8.3, p = .02). Corvids stayed away equally long when chased by either method (χ2(2, N =176) = 2.6, p = 0.3).

## 4. Discussion

There is a need for novel methods to deter birds, and we show that the RobotFalcon can make a major contribution to filling that niche. It cleared fields from corvids, gulls, starlings and lapwings successfully and fast, with deterred flocks staying away for hours. The RobotFalcon was more effective than a drone: its success was higher, and it deterred flocks faster. The effectiveness of the RobotFalcon was similar across flocks of different species, while that of the drone was lower for flocks of gulls and corvids than for starlings. The effectiveness of the RobotFalcon was higher when it approached flocks from a higher altitude, as evidenced by the longer flight initiation distances. This may be because if a RobotFalcon approaches from above it represents a greater potential threat, or because it is detected earlier by the flock.

We compared the RobotFalcon against the most effective methods used at airbase Leeuwarden: distress calls and pyrotechnics. The RobotFalcon kept away flocks of gulls, lapwings and starlings (but not corvids) for longer than the best methods at airbase Leeuwarden. Fields were kept free from corvids equally long when their flocks were deterred by the RobotFalcon and the best airbase methods. This may because corvids depend on the local area for their resources more than gulls, lapwings and starlings. A limitation of our approach is that we compared different methods at different sites (RobotFalcon in Workum versus best airbase methods in Leeuwarden). This comparison may be conservative however, because even though the habitat management by Airbase Leeuwarden made their area less attractive to birds, flocks still returned to them sooner than at fields in Workum.

Effectiveness of most of the current methods to drive away birds is reduced by habituation, with birds fleeing less over time. Birds habituate in particular to methods that do not represent a natural threat (such as synthetic sounds, gas cannons, reflectors and scarecrows), especially when such methods are the only ones used in the field (BSCE, 1988; EIFAC, 1988; Coniff, 1991; Davis & Harris, 1998; Matyjasiak, 2008). The Dutch Air Force resolves this by alternating between different methods (species specific distress calls of birds and pyrotechnics). This alternation prevents habituation, but birds still return sooner than when chased away by the RobotFalcon. In our study, there was no sign of habituation to the RobotFalcon in our three months of field work. We speculate that the continued effectiveness of the RobotFalcon was due to its natural appearance and behaviour. This may also explain why it performed better than the drone. We note however that as we did not mark birds individually, the lack of habituation we recorded can be either caused by us deterring naieve birds each day due to the turnover of the bird population, or it may really reflect a lack of habituation of individual birds. We cannot formally distinguish between these options, but anticipate both processes to have contributed, and emphasize that for practical purposes the salient finding is that there was no discernible decrease in success in clearing fields over the three months of field work. For measuring actual levels of habituation to the RobotFalcon specific experiments in semi-controlled conditions in truly resident bird populations such as domestic pigeons should be carried out.

While the RobotFalcon has proven to be a highly effective tool to deter birds, it is important to also recognize its limitations, which are that steering the RobotFalcon requires trained pilots, flights are limited by battery life (15 minutes per battery) and cannot be conducted in rain. Further, to deter large birds (e.g. geese), a robot that mimicks a natural (larger) predator of large birds is needed (e.g. a RobotEagle). We propose nevertheless that deterrence with the RobotFalcon can replace falconry, because it has the same advantages but without the limitations of live birds of prey. More generally, since the RobotFalcon effectively deters flocks of a large range of bird species, without any signs of habituation, we propose it to be a game-changing addition to the tool-box currently available.

## Author Contributions

**Rolf F. Storms:** Methodology, Investigation, Formal analysis, Data curation, Writing-Original draft preparation. **Claudio Carere:** Writing- Reviewing and Editing. **Robert Musters:** Methodology. **Hans van Gasteren:** Resources, Writing- Reviewing and Editing. **Simon Verhulst:** Writing- Reviewing and Editing. **Charlotte K. Hemelrijk:** Conceptualization, Supervision, Writing- Reviewing and Editing.

## Acknowledgments

Peet Sterkenburgh provided us with the the permissions to deter birds from specific areas in Workum. Ronja Hulst, Sorscha Passmore and Deborah Salleh contributed to the field work. Ramon Wind (RW) was our second certified pilot for the RobotFalcon. Martin Das and Minne Hellinga deterrered birds at airbase Leeuwarden. This publication is part of the project *Preventing bird strikes: Developing RoboFalcons to deter bird flocks* (with project number 14723) of the Open Technology programme which is financed by the Dutch Research Council (NWO) awarded to CKH.

## Data availability statement

Data will be available from the 4TU Reasearch Data Repository upon publication.

