## Supplementary for "Improved deterrence of birds using an artificial predator, the RobotFalcon"

**Corresponding author**

Rolf F. Storms, MSc

Groningen Institute for Evolutionary Life Sciences, University of Groningen

Nijenborg 7

9747 AG Groningen, The Netherlands

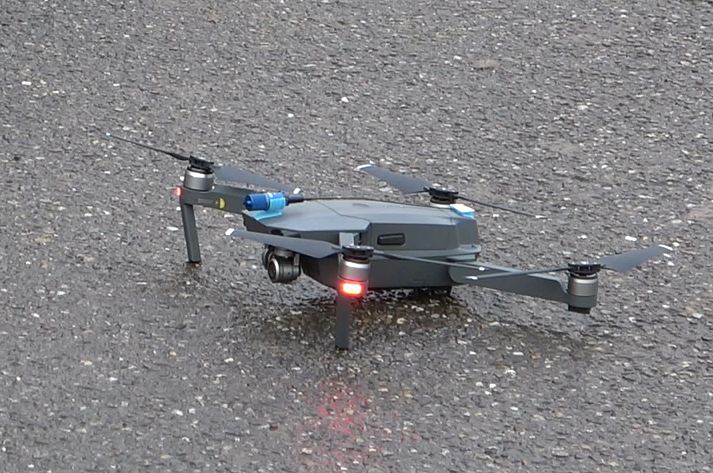

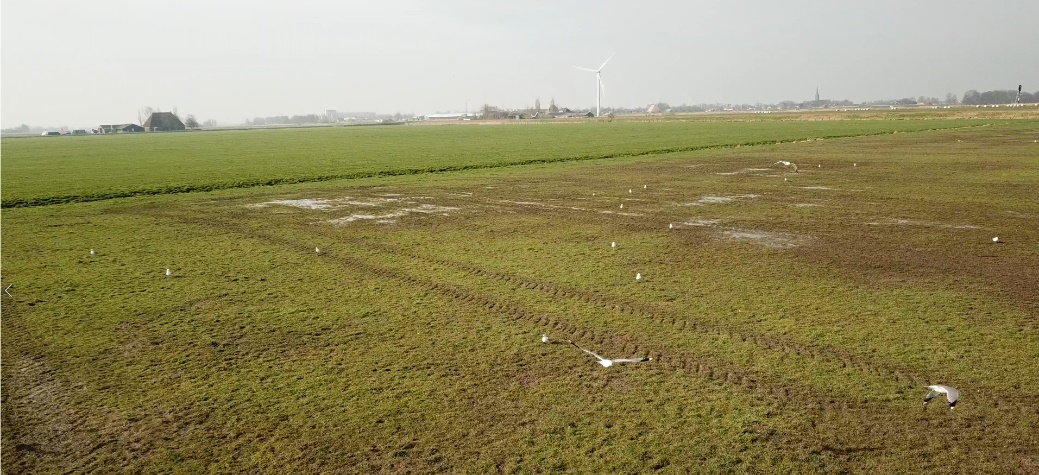

**Figure s1.** The drone (left) and an example of its view during flight (right).

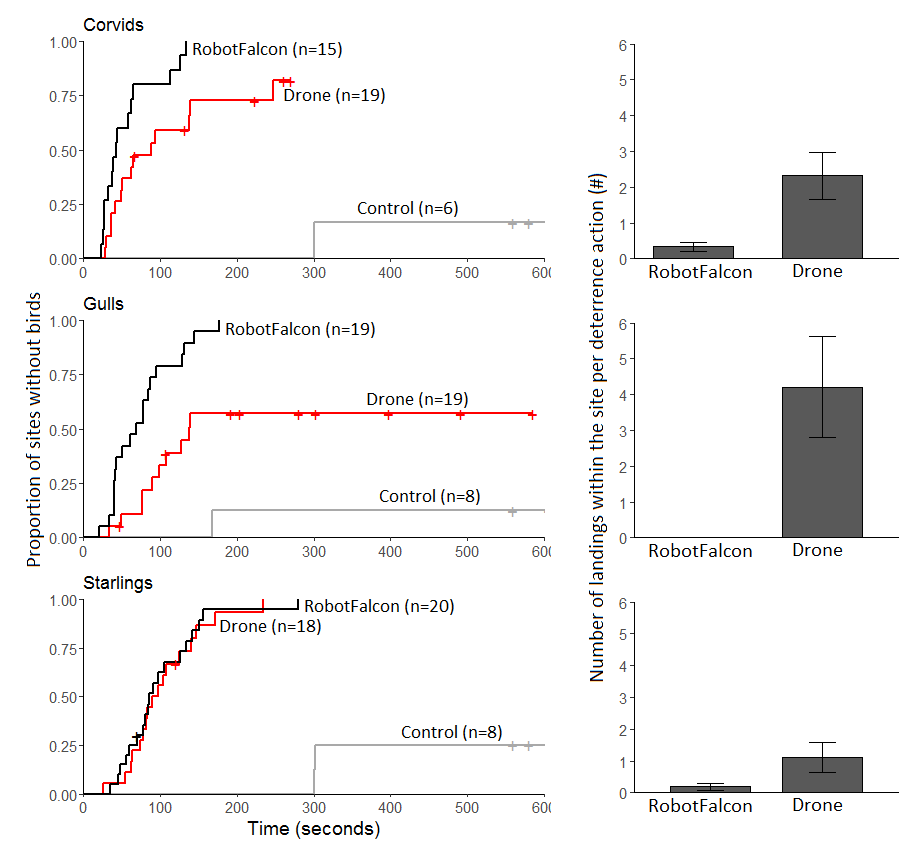
**Figure s2**. Comparison between deterrence of flocks per species with RobotFalcon, drone and control (=no disturbance). (A). Proportion of fields cleared from flocks of birds after deterrence with a RobotFalcon or a drone over time. The RobotFalcon chased away flocks of corvids and gulls significantly faster than the drone (Figure s2, χ2(1, N = 34) = 7.3, p = .007; χ2(1, N = 38) = 12.7, p < .001), but starlings were chased away by both methods equally fast (Figure s2, χ2(1, N = 38) = 0, p = 0.9). (B).The number of times birds landed again within the field during a deterrence action with the RobotFalcon and drone.

**A	B**
